# A new molecular marker including parts of conservative histone H3 and H4 genes and spacer between them for phylogenetic studies in dragonflies (Insecta, Odonata), extendable to other insects

**DOI:** 10.1101/2024.10.28.620759

**Authors:** Anatoly V. Mglinets, Vera S Bulgakova, Oleg E Kosterin

**Affiliations:** Institute of Cytology and Genetics of the Siberian Branch of Russian Academy of Sciences, Acad. Lavrentyev ave. 10, Novosibirsk, 630090, Russia

**Keywords:** histone repeat, histone H3, histone H4, intergenic spacer, Odonata, insects, molecular marker

## Abstract

A new molecular marker, the histone H3-H4 region, containing partial coding sequences of the genes of histones H3 and H4 and the non-coding spacer between them, is proposed. This marker is potentially useful for molecular phylogenetic studies at generic, species, and even intra-species level in insects. The highly conservative histone coding sequences ensure universality of primers and the ease of primary alignment, while the highly variable non-coding spacer provides enough variation for analyses at short evolutionary distances. In insects, the histone genes reside in the histone repeat which is tandemly repeated in dozens to hundred copies forming the so-called histone cluster. However, the order and orientation of the histone genes in the histone repeat is variable among orders, which exerts some limitation for the use of the proposed marker. The marker efficacy is hereby shown for Odonata (dragonflies and damselflies), where it well resolved families, genera and species involved and provided an insight into the relationship of two *Sympetrum* species, *Sympetrum croceolum* and *S. uniforme*. The same combination of original primers should work also in Diptera.

## Introduction

Analysis of DNA variation is a powerful tool in reconstructing phylogenetic history of living creatures, hence the use of molecular methods provided a profound progress in phylogenetic analysis, with applications in taxonomy, paleobiology, paleogeography and evolutionary theory. Particular sequences used for this purpose, traditionally called ‘phylogenetic markers’, differ in their rate of fixation of mutations thus permitting phylogenetic resolution at different time scales, with resolution of most recent divergences being possible with most variable markers, for those applied to Odonata see Cheng et al. (2018).

Mitochondrial DNA in animals is on average more variable than nuclear DNA, so the popular mitochondrial marker cytochrome oxydase I (*COI*) is widely used for barcoding of animals (Ballard & Whitlock 2004; Avise 2009). Howeverm, the phylogenies reconstructed from mitochondrial markers often contradict to both phylogenies reconstructed from nuclear markers and traditional taxonomy based on morphology. In some groups of organisms (Cheng et al., 2023), including the family Coenagrionidae in Odonata (Dow et al. 2019, Deng et al. 2021; Galimberti et al. 2021, Geiger et al. 2021), mitochondria seem to ‘live the life of their own’. Most drastic patterns were revealed in *Coenagrion* Kirby, 1890, where mitochondrial haplotypes cross species barriers but seem unable to cross the Hybraltar Strait (Ferreira et al. 2016; Galimberti et al. 2021, Geiger et al. 2021), and in *Ischnura elegans* (Vander Linden, 1820), where Japanese population appeared strongly divergent from the continental ones (Deng et al. 2021). Such drastic patterns can hardly be explained by incomplete lineage sorting or introgressive hybridisation, the phenomena usually supposed to explain such cases. Their reason could be co-selection of mitochondria with strains of the intra-cellular bacterial endosymbiont *Wolbachia* Hertig, 1826 (Deng et al. 2021). Horizontal transfer of mitochondria via an unknown agent is also postulated but not proved (Gurdon et al. 2016). Another problem of using markers based on mitochondrial DNA are sporadic occurrence of mtDNA fragments adopted by the nuclear genome (NUMT), which are variably divergent from the actual mtDNA and may result in false phylogenetic results; for examples in Odonata see Ožana et al. (2022) or Lorenso-Carballa et al. (2022). For the purpose of revealing evolutionary history at short time distances, microsatellite or SSR markers were for a long time popular and applied also to Odonata (e.g. Lowe et al. 2008). These are tracts of very short (one to few nucleotides) repeats with the number of copies frequently changing because of slippage mispairing and unequal crossing over. Such markers suffer from high rate of homoplasy, when different events of copy number change result in the same alleles, here understood as particular numbers of repeats but still are the marker of choice for population genetic studies.

The modern next generation high throughput sequencing and genomic approach offer ample phylogenetic data (for examples in Odonata see e.g. Futahashi et al., 2015; Bybee et al., 2021; Kohli et al., 2021), potentially useful for analysis even at short evolutionary distances, but are expensive and more demanding to sample preparation. Although the prices are getting lower, these technologies still remain unnafordable for many researchers in countries which harbour the richest biodiversity. Therefore, there is still a need of easily analysed and cheap nuclear markers based on Sanger sequencing which would provide good resolution at the species level and could be useful at least for fast preliminary revealing the evolutionary history of populations, subspecies and closely related species.

Non-coding sequences, whose variability is mainly determined by physical properties of DNA replication, are useful candidates for variable phylogenetic markers. Their use is limited by possibility to work out universal primers that could be achieved by involvement of bordering conserved sequences. Of such markers, the so-called ITS region including internal spacers between the conserved ribosomal RNA genes, *ITS1* and *ITS2,* is the most popular among nuclear markers; for its use in Odonata see Hovmöller & Johansson (2004), Dumont et al. (2010), Karube et al. (2011), Schneider et al. (2023). The highly repetitive nature of the nucleolus organiser provides an advantage of high concentration of the template in preparations of genomic DNA and a disadvantage of possible heterozygosity as well as cis-heterogeneity between individual repeat copies (Hovmöller & Johansson 2004). Recently a useful approach has been proposed, focusing on introns of nuclear genes (Ferreira et al., 2014). The primary structure of introns has scarce adaptive constraints save mutations affecting splicing. At the same time, the bordering exons are usually conservative enough to allow for universal primers design. These markers have a disadvantage of low concentration of template genomic DNA, since the genes involved are unique and present in the genome only in two copies, so they may be less readily amplifiable from specimens with somewhat degraded DNA as compared to the highly repeated sequences of the ITS region.

In the present work we propose and test usability of a new phylogenetic marker, resembling the spacer between ribosomal RNA genes in being a spacer between coding genes and a part of a tandem repeat. This is a spacer between the genes of the conservative core histones H3 and H4 and partial coding sequences of these histones, which we designate as *the histone H3-H4 region*.

In animals, the genes for five histones (H1, H2A, H2B, H3, H4) are included into the so-called histone repeat, tandemly repeated copies of which form the histone cluster (Eirín-López et al., 2009). Several important circumstances should be noted in this respect:

i. Histones H3 and H4 are among the most conserved proteins in eukaryotes (Stein et al. 1984; Doenecke et al. 1997; Eirín-López et al. 2009), so their coding sequences allow primers of very broad applicability across taxonomic groups.
ii. In insects, genes of these two histones are disposed in the histone repeat relatively close to each other (Eirín-López et al. 2009). The order of histone genes and orientation of their reading frames is variable among different insect orders. For instance, their order is H1, H3, H4, H2A, H2B, in *Drosophila melanogaster* (Goodenough 1984: p. 304; Eirín-López et al. 2009) and in all Odonata species tested in this work (Fig. 1). Although this variation may also take place at lower taxonomical levels and is sometimes observed even in different copies of the histone repeat in the same chromosome, in all species of Odonata tested by us (see below) we obtained a PCR product using the same primer pair matching histones H3 and H4. So the original primers we suggest are useful at least across dragonflies and damselflies.
iii. The spacer between H3 and H4 histone genes is non-coding and therefore is expected to undergo neutral evolution, hence being a kind of molecular clock.
iv. (iv). Insects have hundreds of copies of the histone repeat (Solovyev et al. 2022), that facilitates amplification from total genomic DNA preparations, but may also bring about problems related to cis-(within-cluster) and trans-(allelic between homologous chromosomes) heterogeneity..
v. Most insects have only one histone cluster, while in some of them and in other animal groups there is a number of paralogous clusters (Eirín-López et al. 2009). A single histone cluster is an advantage since this avoids trans-cluster heterogeneity which would complicate an analysis.

**Figure 1.**
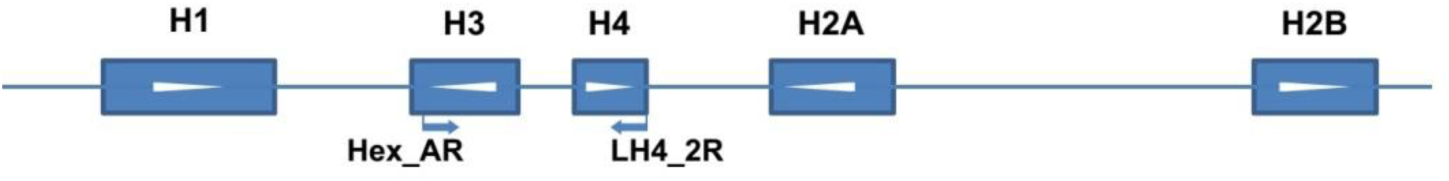
Positions of the primers used to amplify the histone H3-H4 region in the histone repeat as exemplified by a fragment of assembly of *Ischnura elegans* genome (NW_025791746). White arrowheads indicate direction of transcription.

To develop and test the marker we chose the order Odonata and designed original primers which worked in all tested species. We tested its resolution at different taxonomical levels, by sequencing amplicons from representatives of different families (Calopterygidae, Coenagrionidae, Aeshnidae, Gomphidae, Corduliidae and Libellulidae), from several species of some genera and from a number of specimens of some *Sympetrum* spp. The latter involved series of three species from the same *danae* species group (Pilgrim et al. 2012), namely *Sympetrum croceolum* (Selys, 1883), *S. danae* (Sulzer, 1776) and *S. uniforme* (Selys, 1883), including those collected in the same populations.

## Materials and methods

### Material

The specimens of *Sympetrum croceolum* (Selys, 1883)*, S. danae* (Sulzer, 1776)*, S. flaveolum* (Linnaeus, 1758), and *S. uniforme* (Selys, 1883) were preserved in 96% ethanol, other specimens were treated overnight with acetone and then dried out. The species and specimens from which the histone H3-H4 region was sequenced in the course of this study and the GenBank accession numbers of these sequences are enumerated (in parentheses) in Table 1.

**Table 1.** Species (by families) and specimens sequenced for the histone H3-H4 region, their origin and the GenBank accession numbers of the sequences. Coordinates are given in decimal degree format.

| Species name | locality | latitude (N) | longitude (E) | date | specimens, their codes (if any) and accession numbers of their sequences |
| --- | --- | --- | --- | --- | --- |
| Calopterygidae |  |  |  |  |  |
| *Atrocalopteryx atrata* (Selys, 1853) | Russia, Khabarovskiy Kray, 44 km SE of Khabarovsk, Chirki, | 48.19 | 134.68 | 15.07.2014 | 1 ♂ (PQ498535) |
| Coenagrionidae |  |  |  |  |  |
| *Coenagrion johanssoni* (Wallengren, 1894) | Russia, Khabarovskiy Kray, 44 km SE of Khabarovsk, Chirki | 48.19 | 134.68 | 15.07.2014 | 1 ♂ (PQ498539) |
| Aeshnidae |  |  |  |  |  |
| *Aeshna juncea* (Linnaeus, 1758) | Russia, Khabarovskiy Kray, 28 km SE of Khabarovsk, Bychikha village | 48.30 | 134.83 | 9.07.2014 | 1 ♂ (PQ498536) |
| Gomphidae |  |  |  |  |  |
| *Nihonogomphus raptus* (Selys in | Russia, Khabarovskiy Kray, 44 km SE of Khabarovsk, Chirki ranger | 48.19 | 134.68 | 15.07.2014 | 1 ♂ (PQ498542) |
| Hagen, 1858) | station |  |  |  |  |
| Corduliidae |  |  |  |  |  |
| *Cordulia aenea*<br>(Linnaeus, 1758) | Russia, Novosibirsk Academy<br>Town | 54.8473 | 83.1072 | 1.07.2015 | 1 ♂ (PQ498537) |
| *Somatochlora*<br>*arctica* (Zetterstedt,<br>1840) | Russia, Novosibirsk Academy<br>Town, Zvezda Beach | 54.8314 | 83.0538 | 30.06.2015 | 1 ♀ (PQ498546) |
| *Somatochlora*<br>*graeseri* Selys, 1887 | Russia, Primorskiy Kray, Khanka<br>District, Lake Khanka W bank 2<br>km N of Platono-Aleksandrovskoe<br>village | 45.062 | 131.995 | 4.07.2014 | 1 ♀ (PQ498561) |
| Macromiidae |  |  |  |  |  |
| *Macromia*<br>*amphigena fraenata*<br>Martin, 1906 | Russia, Khabarovskiy Kray, 44 km<br>SE of Khabarovsk, Chirki | 48.19 | 134.68 | 15.07.2014 | 1 ♂ (PQ498540) |
| Synthemistidae *sensu lato* |  |  |  |  |  |
| *Macromidia rapida*<br>Martin, 1907 | Cambodia, Monduliri Province, a<br>left tributary of the Monorom<br>Waterfall river, 3.5 km SE of Sen<br>Monorom | 12.441 | 107.159 | 13.06.2014 | 1 ♂ (PQ498541) |
| Cordulegastridae |  |  |  |  |  |
| *Cordulegaster picta*<br>Selys, 1854 | Russia, Krasnodarskiy Kray,<br>Kabardinka village, the Krasnaya<br>Shchel' Valley | 44.671 | 37.918 | 30.07.2015 | 1 ♀ (PQ498538) |
| Libellulidae |  |  |  |  |  |
| *Orthetrum albistylum*<br>(Selys, 1848) | Russia, Primorskiy Kray, Khanka<br>District, Lake Khanka W bank 2<br>km N of Platono-Aleksandrovskoe<br>village | 45.062 | 131.995 | 4.07.2014 | 1 ♀ (PQ498543) |
| *Orthetrum glaucum*<br>(Brauer, 1865) | Cambodia, Monduliri Province, a<br>brook downstream Buu Sraa<br>Waterfall | 12.570 | 107.418 | 10.06.2014 | 1 ♂ (PQ498544) |
| *Rhyothemis phyllis*<br>(Sulzer, 1776) | Cambodia, Kampot Province, a left<br>oxbow of the Kampot River<br>downstream of Tek Chhou Rapids | 10.672 | 104.137 | 21.08.2011 | 1 ♀ (PQ498545) |
| *Sympetrum*<br>*arenicolor* Jödicke,<br>1994 | Iran, Lorestan, Khorramabad<br>County, 3.2 km NNW Pasil village | 33.33 | 48.88 | 26.05.2017 | 1 ♂ (PQ498547) |
| *Sympetrum*<br>*cordulegaster* | Russia, Khabarovskiy Kray, 28 km<br>SE of Khabarovsk, a pond S of<br>Bychikha village | 48.292 | 134.829 | 21-22.08.2016 | 1 ♂ (PQ498588) |
| *Sympetrum*<br>*croceolum* (Selys,<br>1883) | Russia, Amursk District,<br>Bolon'skiy Nature Reserve, Kirpu<br>ranger station | 49.506 | 136.028 | 15.09.2016 | 10 ♂♂ (Sc-1 –<br>Sc-10;<br>PQ498574–<br>PQ498582) |
| - “ - | Russia, Khabarovskiy Kray, 28 km<br>SE of Khabarovsk, Bychikha<br>village | 48.292 | 134.829 | 21-22.08.2016 | 2 ♀♀ (Sc-31, Sc-32; Q498583,<br>PQ498586) |
| - “ - | Russia, Altai Republic, Mayma<br>District, Lake Manzherok SE bank | 51.82 | 85.81 | 16.09.2016 | 2 ♂♂ (Sc-41, Sc-42; PQ498584, |
|  |  |  |  |  | PQ498585) |
| *Sympetrum danae*<br>(Sulzer, 1776) | Russia, Novosibirsk, Academy Town, botanical garden | 54.825 | 83.114 | 8.10.2015 | 12 ♂♂ (Sd-1, Sd-3 – Sd-12; PQ498549–PQ498557) |
| *Sympetrum flaveolum* (Linnaeus, 1758) | Russia, Novosibirsk, Academy Town, botanical garden | 54.825 | 83.114 | 8.10.2015 | 1 ♂ (Sf-1; PQ498558), 1 ♀ (Sf-2; PQ498559) |
| - “ - | Russia, Primorskiy Kray, Lake Khanka W bank just N of Platono-Aleksandrovscoe village env | 45.06 | 131.99 | 4.07.2014 | 1 ♀ (Sf-3; PQ498560) |
| *Sympetrum fonscolombii* (Selys, 1840) | Iran, Esfahan Province, Golpayegan City, the Ghomrood (Anaarbar) River | 33.46 | 50.26 | 21.05.2017 | ♂ (No. 1) |
| - “ - | Russia, Krasnodarskiy Kray, Bol’shoi Utrish village env. | 44.76 | 37.39 | 5.08.2015 | 1 ♂ (No. 2; PQ498589) |
| *Sympetrum pedemontanum* (Müller in Allioni, 1766) | Russia, Novosibirsk, a glade in a pine forest between Nizhnyaya El’tsovka and Pravye Chyomu estates | 54.864 | 83.051 | 4.08.2018 | 1 ♂ (PQ498591) |
| *Sympetrum sanguineum* (Müller, 1764) | Russia, Novosibirsk Province, Kyshtovka District, a roadside swamp in the Kyshtovka village NW suburbs | 56.579 | 76.603 | 14.07.2015 | 1 ♂ (Ss-1, PQ498590) |
| - “ - | Krasnodarskiy Kray, Lake Krugloe | 44.677 | 37.594 | 10.06.2016 | 1 ♂ (Ss-2) |
| *Sympetrum striolatum* (Charpentier, 1840) | Iran, Lorestan, Borujerd County, Chenar Chashmah | 33.798 | 40.905 | 25.05.2017 | 1 ♂ (PQ498592) |
| *Sympetrum vulgatum* (Linnaeus, 1758) | Russia, Krasnodarskiy Kray, Kabardinka village env., Krasnaya Shchel’ Valley | 44.672 | 37.922 | 6.07.2016 | 1 ♂ (PQ498593) |
| *Sympetrum uniforme* (Selys, 1883) | Russia, Khabarovskiy Kray, 28 km SE of Khabarovsk, Bychikha village | 48.292 | 134.829 | 21-22.08.2016 | 5 ♂♂, 4 ♀♀ (Su-21–Su-29; PQ498565–PQ498573) |
| - “ - | Russia, Khabarovskiy Kray, Amursk District, Bolon’skiy Nature Reserve, Kirpu ranger station | 49.506 | 136.028 | 15.09.2016 | 1 ♂ (Su-1; PQ498563), 1 ♀ (Su-2; PQ498564) |

### Sequences from public databases

To expand our sample, we downloaded sequences of the H3-H4 region from the Whole Genome Sequence datasets available in public databases of 16 more Odonata species: *Hetaerina americana* (Fabricius, 1798), *H. titia* (Drury, 1773) (Calopterygidae), *Argia fumipennis* (Burmeister, 1839), *Ceriagrion tenellum* (De Villers, 1789), *Ischnura elegans*, *I. senegalensis* (Rambur, 1842) *Pseudagrion microcephalum* (Rambur, 1842), *Pyrrhosoma nymphula* (Sulzer, 1776) (Coenagrionidae), *Platycnemis pennipes* (Pallas, 1771), *Prodasineura notostigma* (Selys, 1860) (Platycnemididae), *Tanypteryx hageni* (Selys, 1879), *Tachopteryx thoreyi* (Selys, 1889), *Uropetala carovei* (White in Dieffenbach, 1843) (Petaluridae), *Brachytron pratense* (Müller, 1764) (Aeshnidae), *Pachydiplax longipennis* (Burmeister, 1839), *Pantala flavescens* (Fabricius, 1798), (Libellulidae).

Besides, the histone H3-H4 region was assembled from SRA archives available atGenBank, with the use of MIRA software (Chevreux et al. 1999) for the following 20 species: *Archilestes grandis* (Rambur, 1842) (Lestidae), *Calopteryx splendens* (Harris, 1780), *Hetaerina vulnerata* Hagen in Selys, 1853, *Mnais tenuis* Oguma, 1913, *Neurobasis kaupi* Brauer, 1867 (Calopterygidae), *Agriocnemis femina* (Brauer, 1868) (Coenagrionidae), *Anax parthenope* (Selys, 1839), *A. strenuus* Hagen, 1867 (Aeshnidae), *Gomphus vulgatissimus* (Linnaeus, 1758), *Lanthus parvulus* (Selys, 1854), *Onychogomphus forcipatus* (Linnaeus, 1758), *Ophiogomphus mainensis* Packard in Walsh, 1863 (Gomphidae), *Cordulegaster boltonii* (Donovan, 1807) (Cordulegastridae), *Macromia manchurica* Asahina, 1964 (Macromiiddae), *Ladona fulva* (Müller, 1764), *Leucorrhinia albifrons* (Burmeister, 1839), *Libellula angelina* Selys, 1883*, L. quadrimaculata* (Linnaeus, 1758), *Nannophya pygmaea* Rambur, 1842, *Orthetrum coerulescens* (Fabricius, 1798) (Libellulidae). The accession numbers of the database entries used as sources of these sequences are indicated at species names in Figs 2-3.

**Figure 2.**
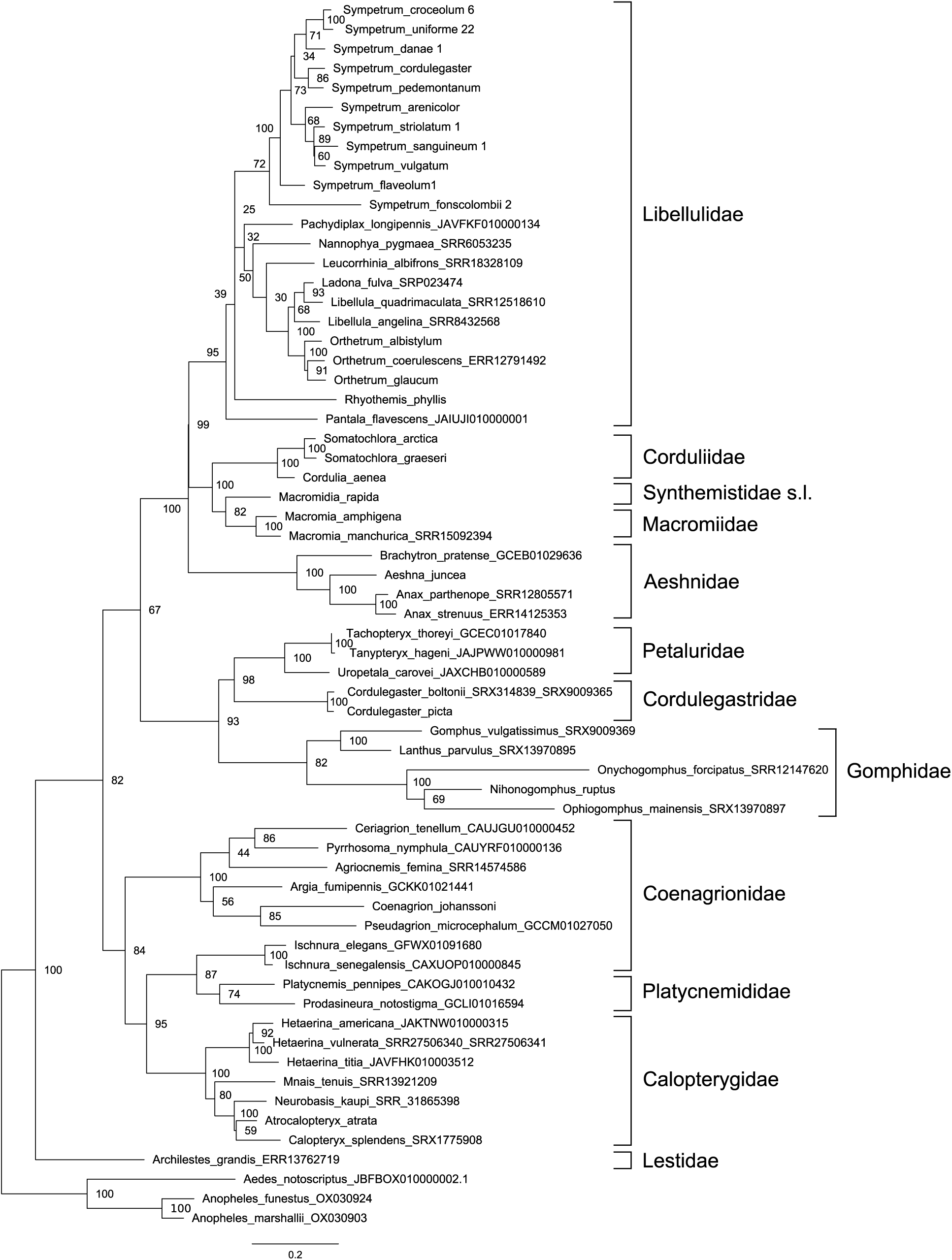
Phylogenetic tree of the studied species of Odonata reconstructed with Maximum Likelihood method from the histone H3-H4 region sequences. Boostrap values are shown at respective nodes. Three species of Diptera, Culicidae serve as the outgroup.

**Figure 3.**
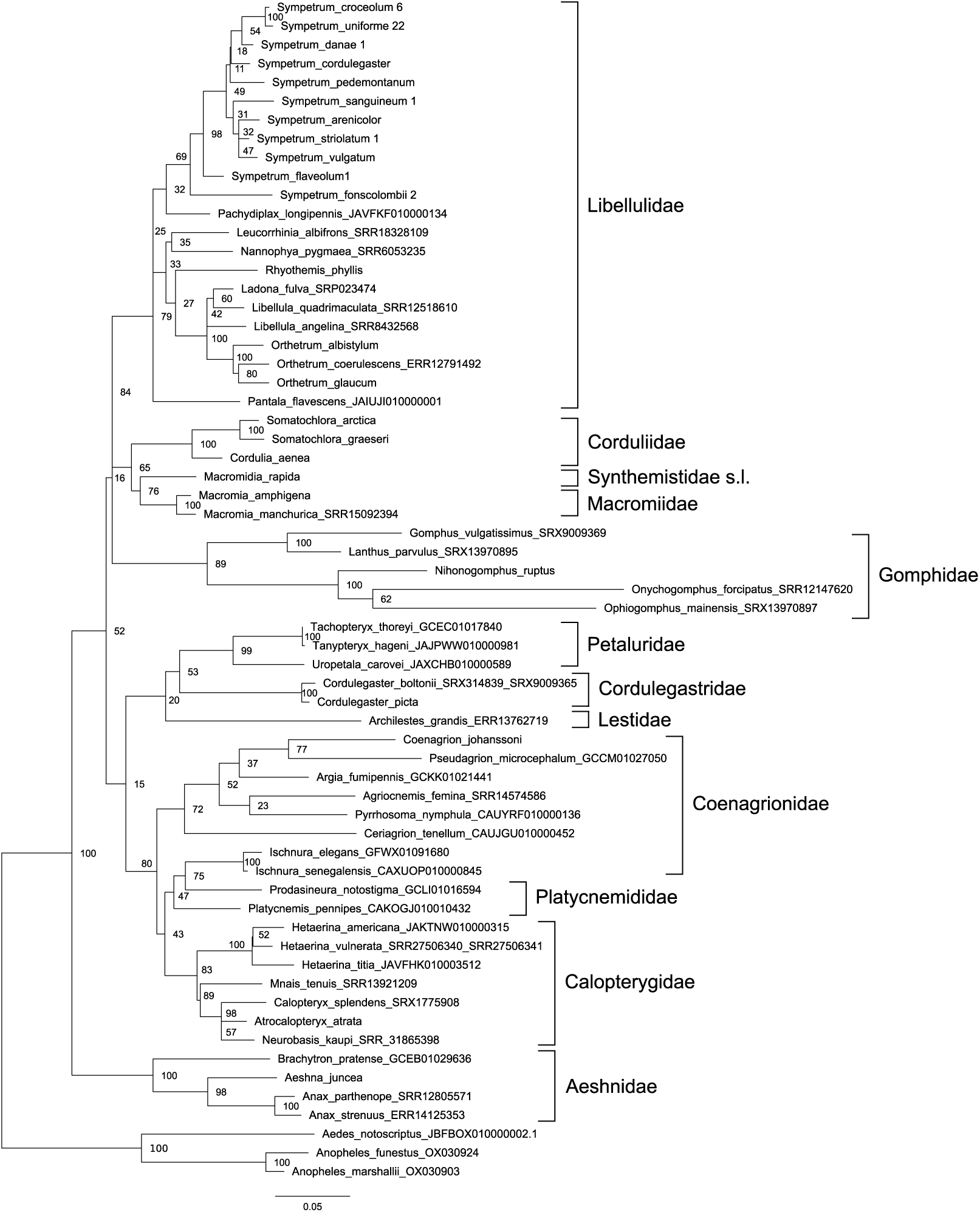
Phylogenetic tree of the studied Odonata species reconstructed with Maximum Likelihood method from the fragments of coding sequences of histone H3 and histone H4 genes involved into the proposed ‘histone H3-H4 region’ marker. Boostrap values are shown at respective nodes.

### Primer design

We designed some 14 original primers to match different parts of insect histone genes coding for H1, H2B, H3, H4 comprising the histone gene cluster. At the start of the present work we did not know the precise order and orientation of the histone genes in the cluster, so we tested different primer combinations to select primer pair(s) which would produce an amplification product containing the fragments of the genes of histone H3 and H4 and a spacer between them. We found out that the pair of primers Hex_AR matching the 3’ portion of the H3 gene (in the orientation opposite to that of transcription) and LH4_2R matching the 3’ portion of the H4 gene (also in the orientation opposite to that of transcription) (Fig. 1) produced the expected product, indicating that the H3 and H4 genes were oriented anti-parallel as to their reading frames, with 5’ ends of their coding chains oriented towards each other (Fig. 1). The primer sequences are as follows:

Hex_AR: 5’ atatccttgggcatgatggtgac (forward)

LH4_2R: 5’ ttaaccgccgaaaccgtacagggt (reverse)

The primer LH4_2R, matching the coding region of the histone H4 of the moth *Bombyx mori* (Linnaeus, 1758) (Lepidoptera: Bombycidae) (GenBank accession AADK01010708), was worked out in the course of our previous study of the variation of the histone H1 gene in some Lepidoptera (Solovyev et al. 2015), although this particular primer was not mentioned in the cited work and is published here for the first time.

The Hex_AR primer was worked out to match the coding sequences of the histone H3 gene of *Ophiogomphus severus* (Hagen, 1874) (Odonata: Gomphidae) taken from GenBank (accession AY125228).

The conding sequences of histones H3 and H4 are well conserved (Stein et al. 1984; Doenecke et al. 1997; Eirín-López et al. 2009) so that the primers worked out to match a particular sequence, one of which was from Odonata and the other from Lepidoptera, worked well for all tested species of Odonata.

### DNA isolation, sequencing and analysis

Dragonfly legs were homogenized in a mortar in 0.2 ml of isolation buffer (0.1 Tris-HCl, pH 8.0; 0.05 M EDTA; 1.25% SDS; 0.5 M NaCl) with Al_2_O_3_ as grinding particles, then mixed with 0.8 ml of the same buffer. The mixture was incubated for 1 hr at 55°C, then added with 350 μl of 5 M potassium acetate, incubated for 30 min on ice and centrifuged at 16.1 g for 10 min. The supernatant was transferred to fresh tubes, mixed with 0.6 ml of isopropanol, incubated at room temperature for 1 hour and centrifuged at 12.2 g for 10 min. The precipitate was washed twice with 0.1 ml 70% ethanol with subsequent centrifugation at 12.2 g for 5 min, dried at 50°C for 5 min and dissolved in 50μl of deionized water.

PCR reaction was carried out in a volume of 20μl with 2 μl of 10х ammonium sulphate buffer, 2 μl of 25 mM MgCl_2_, 0.3 μl of the Hot Start Tаq polymerase produced by SIBENZYME company, Novosibirsk (5 U/μl), 0.15 μl BSA (10 mg/ ml), 1 μl of forward and reverse primers (10 pM) each, 2 μl of 2 mM dNTPs, 2 μl of diluted DNA (20-60 ng) and 10.55 μl of deoinised water. For PCR, BIO-RAD MyCycler thermal cycler was used, with the reaction parameters as follows: denaturation at 95 °C for 3 min followed by 32 cycles including denaturation at 94 °C for 30 s, annealing at 55 °C for 25 s, elongation at 72 °C for 45 s. PCR-products were purified with Invisorb® Spin Filter PCRapid Kit and Sanger sequenced using Big Dye Terminators version 3.0 or 1.1 at SB RAS Genomic Core Facility.

Raw trace files were visualized and translated into nucleotide sequences with the use of the Gap4 software (Staden et al. 2003). The sequences were aligned with ClustalW (Larkin et al., 2007) using MEGA 6.0 software package (Tamura et al. 2013) with default parameters.

The phylogenetic relationships were reconstructed with the Maximum Likelihood method using MEGA 6.0, with Kimura 2-parameter substitution model, as default in the package, rate among sites: gamma-distributed with invariant sites. Bootstrap values from 100 replications were calculated. The sequences of the histone H3-H4 region of three species of Diptera, *Aedes notoscriptus* (Skuse, 1889), *Anopheles funestus* Giles, 1900 and *A. marshalli* Theobald, 1903) from GenBank, were used as the outgroup for the order Odonata-wide phylogenetic reconstruction (Figs 2-4), because in Diptera we found the same order and orientation of the genes of histones H3 and H4.

**Figure 4.**
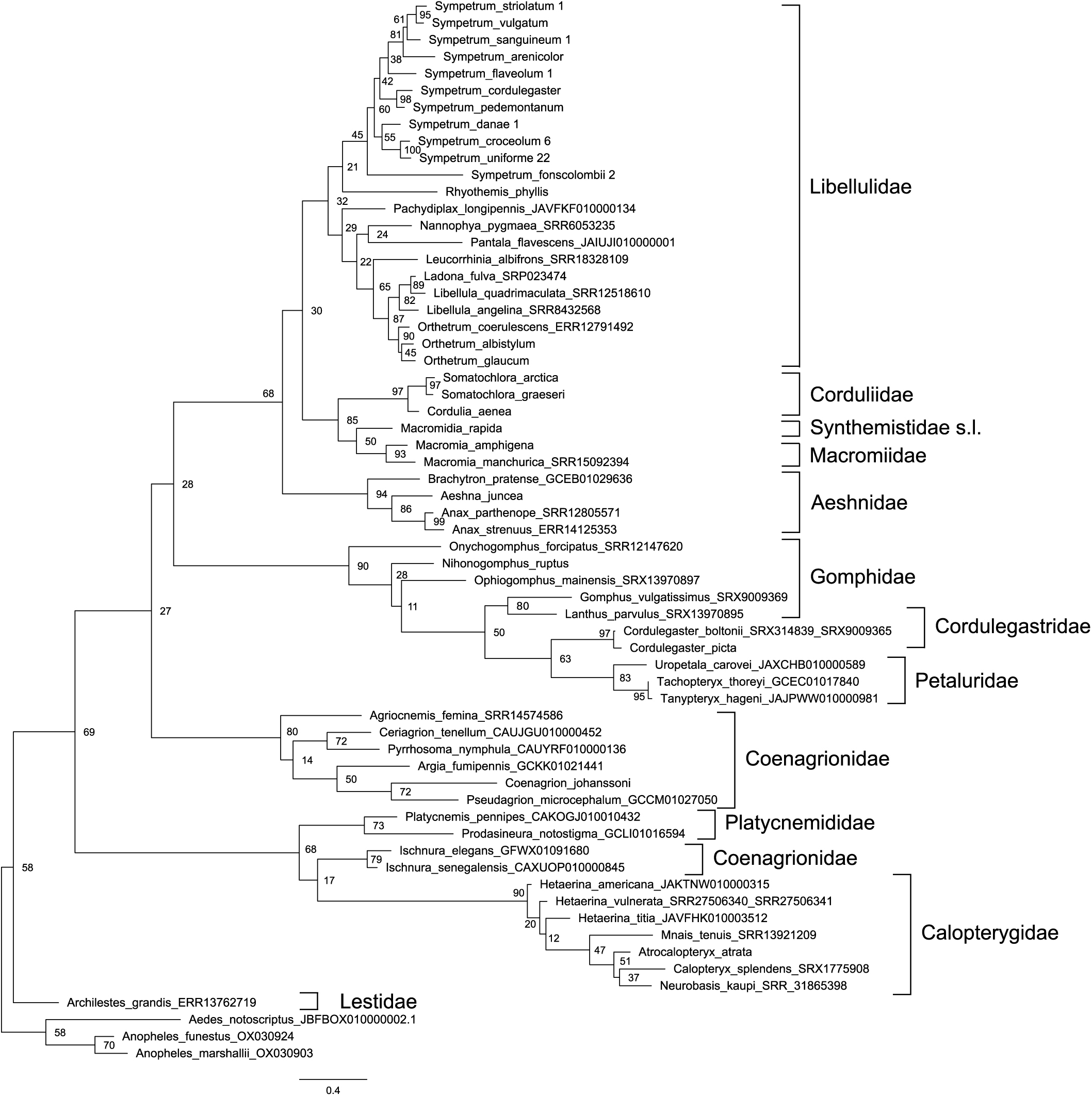
Phylogenetic tree of the studied Odonata species reconstructed with Maximum Likelihood method from the intergenic spacer between histone H3 and histone H4 genes. Boostrap values are shown at respective nodes.

The uncorrected p-distances between different alleles of the histone H3-H4 regions within species of *Sympetrum* were calculated with the MEGA 6.0 software (the entire matrix is not shown).

## Results

The histone H3-H4 region was successfully amplified with the above suggested primer pair and sequenced from DNA isolated from specimens of Odonata enumerated in Table 1, 59 individuals of 24 species. Together with the 36 sequences adopted from public databases, this comprised a sample of 95 sequences of 60 species.

The sequences of the histone H3-H4 region contained parts of the conservative coding sequences of the genes of histones H3 (351 bp) and H4 (288 bp) and the spacer between them of a variable length about 250 bp. All substitutions revealed in the coding sequence fragments were synonymous except for the substitution T˃A in the first position of the second codon of the histone H4 gene, which changes threonine to serine, in both sequenced specimens of *S. fonscolombii* (Selys, 1840) (not shown). As expected, the sequences of bordering coding sequences were unambiguously aligned. At the same time the spacer expectedly appeared hyper-variable so its alignment was much less certain and retained some ambiguity.

One specimen of *S. sanguineum* (Müller, 1764) (Ss-2), one specimen of *S. fonscolombii* (No. 1), and one specimen of *S. uniforme* (Su-23) appeared heterogeneous containing reads with and without deletion of a number of nucleotides in the spacer. One of those indels found in *S. sanguineum* (Ss-2) concerned just one base pair, however, we were able to infer both sequence variants from the chromatogram but used for further analysis only one of them, chosen randomly. Indels found in the other two species were longer, about 5 and 10 bp. Although the sequences beyond the deleted region could be reconstructed, we preferred to exclude these specimens from further analyses.

In some positions, the chromatograms revealed two peaks of comparable height suggesting within-specimen heterogeneity for nucleotides occupying these positions. Those positions reflected either heterozygosity for different alleles or cis-heterogeneity for the histone repeat along a histone cluster, quite expectable in the case of repeated units. Such positions made uncertain the exact number of unique alleles found in a species. Few chromatograms did not resolve nucleotides in a number of positions adjacent to the primers, we nevertheless involved the shortened, well resolved sequences into phylogenetic reconstructions.

First, we reconstructed a phylogenetic tree based on the sequence of the H3-H4 region from one representative of each involved species, both newly sequenced and available or reconstructed from public databases (Fig. 2) The tree was rooted with the sequences of mosquitoes *Anopheles* and *Aedes* used as outgroup. The overall tree topology well corresponded to the family system of Odonata (Dijkstra et al. 2013), except for the odd position of the genus *Ischnura*, which is attributed to Coenagrionidae but clustered with Platycnemididae, although with a weak bootstrap support of 83. Our tree included 10 currently recognised families of Odonata (Dijkstra et al. 2013) represented by several species. Seven of them of them were revealed as monophyletic clades well supported by high bootstrap values: Libellulidae (95), Corduliidae (100), Macromiidae (100), Aeshnidae (100), Petaluridae (100), Cordulegastridae (100) and Calopterygidae (100). Two families appeared monophyletic with weak support: Gomphidae (82) and Platycnemididae (74). The cluster of Coenagrionidae without *Ischnura* had the highest support of 100. Even representatives of the three families (Corduliidae, Macromiidae and Synthemistidae s. l.), previously considered in the family Corduliidae in the broad sense, also grouped in a cluster with the maximum support of 100.

*Archilestes grandis* is the only involved representative of Lestidae, the family considered to retain most plesiomorphic characters among Odonata (Dijkstra et al. 2013). Hence its position as the most basal branch of Odonata was rather expected. If to exclude this branch formally attributed to Zygoptera, both suborders Anisoptera and Zygoptera appeared monophyletic but weakly supported (67 and 84, respectively).

The tree of Fig. 2 includes nine genera represented by more than one species. Seven of them (*Orthetrum* Newmann, 1833, *Somatochlora, Macromia* Rambur, 1842*, Anax* Leah in Breuster, 1815*, Cordulegaster* Leah in Breuster, 1815*, Ischnura* Charentier, 1840*, Hetaerina* Hagen in Selys, 1853) had the highest support of 100. The genus *Sympetrum*, represented by 11 species, formed a monophyletic cluster with a weak support (72). However, if to exclude the problematic (see below) divergent species *S. fonscolombii*, the remained 10 species clustered together with the maximum support of 100. The genus *Libellula* Linnaeus, 1758 would became monophyletic but weakly supported (68) if to assume *Ladona* Needham, 1897 to be its synonym, as is often considered.

It was interesting to evaluate separate inputs into this phylogenetic resolution of the histone coding sequences and spacer, so we reconstructed phylogenetic trees based on these two components separately (Supplementary material, Figs 3 and 4). In both trees, terminal branches uniting close species or genera are mainly well supported. The support of families is somewhat lower than in the tree based on the entire histone H3-H4 region (Fig. 2), with the values in the tree based on the concatenated coding sequences of both histone genes (Fig. 3) being in general higher than in the tree based on the non-coding spacer (Fig. 4). The principal topology of the tree based on the spacer sequences (Fig. 4) remained similar to that of the tree based on the entire H3-H4 region (Fig. 2), but is not supported. The overall topology with respect to positions of families of the tree based on the coding sequences (Fig. 3) is different, does not reflect dichotomy for the two suborders and is even less supported than in the spacer tree (Fig. 4). This can be attributed to saturation of conservative histone gene sequences by synonymous substitutions at long evolutionary distances. Altogether, we may conclude that both parts of the histone H3-H4 region have their input into its resolving power, but the best result is produced by the two parts taken together.

As stated above, the here proposed phylogenetic marker, the histone H3-H4 region, is similar to the popular ITS region containing rRNA genes and two non-coding spacers between them. To compare phylogenetic resolution of these two markers, we reconstructed a phylogenetic tree based on the ITS region sequences adopted from GenBank, which contains the same species except for *N. ruptus* and *S. arenicolor* (Fig. 5). It appeared substantially inferior in resolving odonate families as compared to the H3-H4 tree (Fig. 2). The ITS tree contained a number of awkwardly placed species. The zygopteran *Archilestes grandis* occurred among Anisoptera where it clusters with *U. carovei*, which in turn does not cluster with the two other Petaluridae. *P. pennipes* does not cluster with the second Platycnemididae, *P. notostigma*, but occurs among representatives of Coenagrionidae. The branch of Macromiidae does not cluster with other Anisoptera. *S. fonscolombii* is far decoupled from other Sympetrum spp. We may conclude that the ITS region, is unable to adequately resolve the phylogeny at the level of Odonata families. Boostrap values are shown at respective nodes. Two species of Diptera, Culicidae serve as the outgroup.

**Figure 5.**
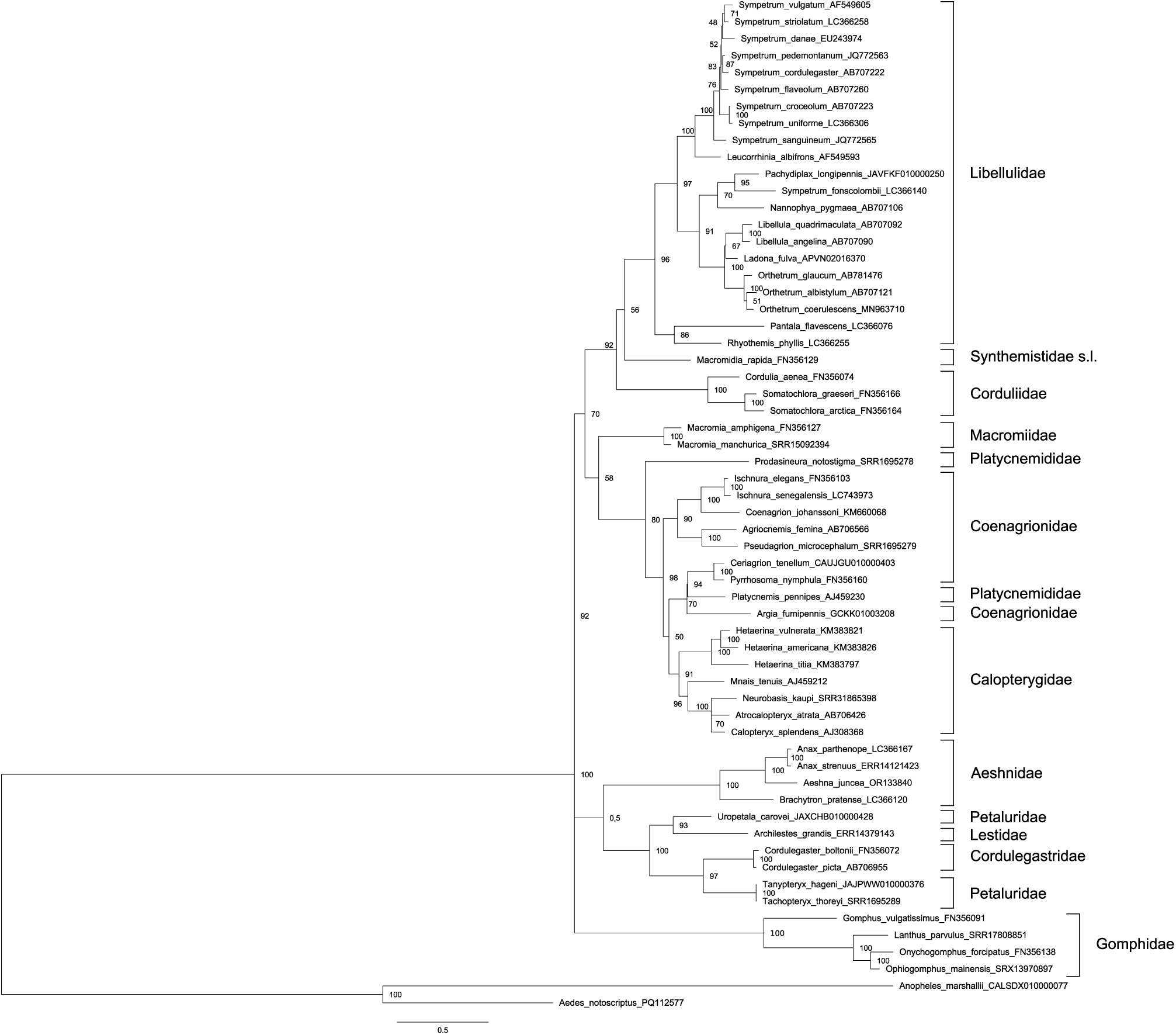
Phylogenetic tree of the species of Odonata as in Fig. 2 (with two omissions) reconstructed with Maximum Likelihood method from the ITS region sequences adopted from GenGank.

To test applicability of the proposed marker, the histone H3-H4 region, to evaluating intra-generic and intra-species variation we estimated its variation and reconstructed a phylogenetic tree for 45 specimens belonging to 11 species of the genus *Sympetrum* involved, using the sequences of *R. phyllis*, *O. albistylum* and *O. glaucum* as the outgroup (Fig. 6). The magnitude of intra-species variation of the histone H3-H4 region sequence appeared quite substantial. For the three species represented by 10 to 14 specimens, *S. croceolum, S. uniforme* and *S. danae*, the maximum uncorrected p-distances (that is, the share of variable positions among all positions) between different alleles within a species appeared to be respectively 0.0216, 0.0037 and 0.0011, that is ca 2.1, 0.4 and 0.1%. The averaged difference between any two sequences within *S. croceolum, S. uniforme* and *S. danae* were 0.0165, 0.0009 and 0.0003, respectively.

**Figure 6.**
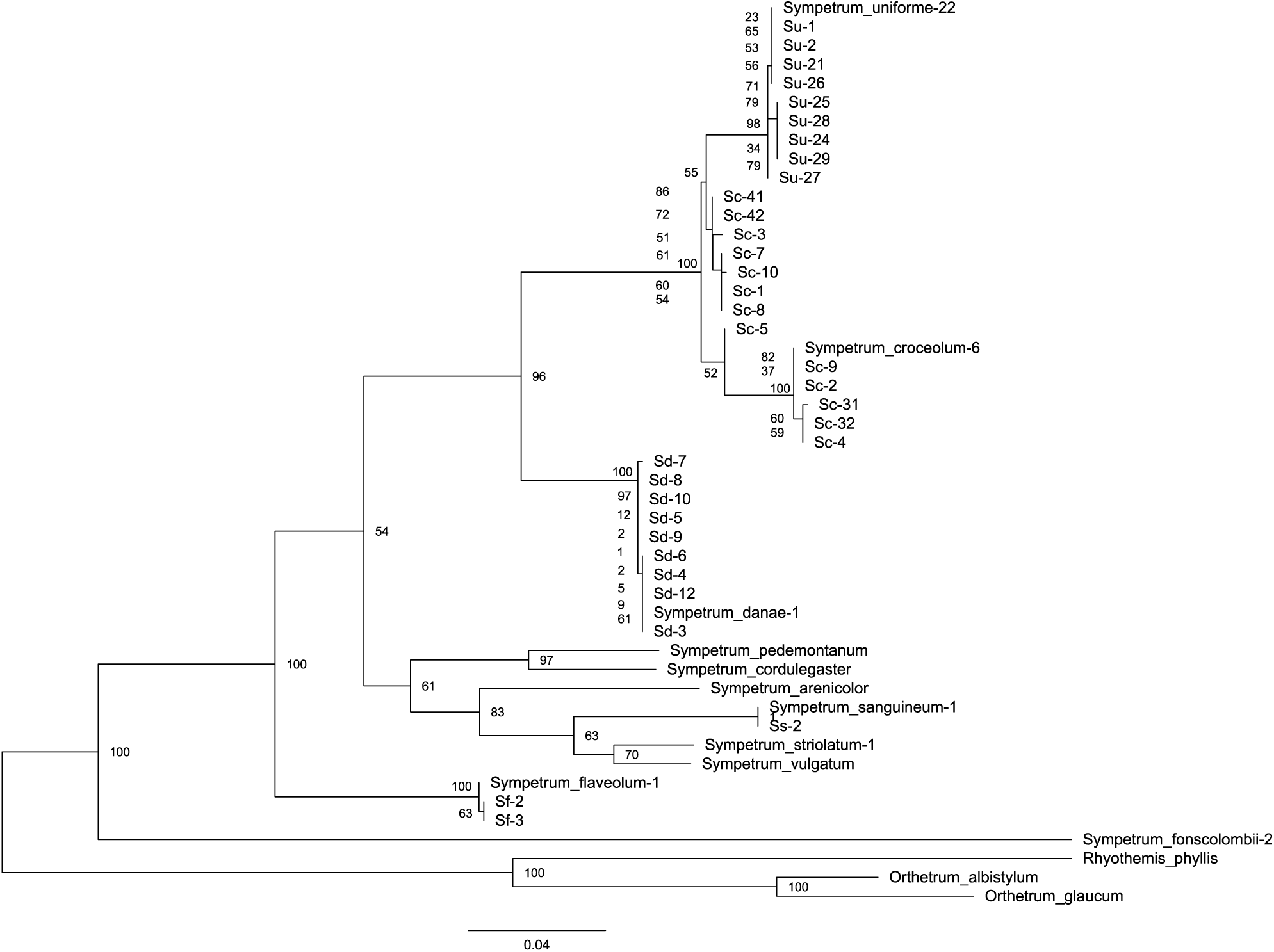
Phylogenetic tree of the studied *Sympetrum* species reconstructed with Maximum Likelihood method from the histone H3-H4 region sequences. Boostrap values are shown at respective nodes. *Orthetrum albistylum* (Selys, 1848), *O. glaucum* (Brauer, 1865) and *Rhyothemis phyllis* (Sulzer, 1776) serve as outgroup.

In the reconstructed phylogenetic tree (Fig. 6), ten sequences of *S. uniforme*, ten sequences of *S. danae* and three sequences of *S. flaveolum* expectedly clustered with the maximum bootstrap support of 100. Strikingly, the cluster of *S. uniforme* appeared to be nested inside that of *S. croceolum*, with the united cluster of these two species also having the support of 100. *S. pedemontanum* (Müller in Allioni, 1766) and *S. cordulegaster* clustered with a support of 76. At the same time, *S. fonscolombii* showed a very deep divergence from the rest of *Sympetrum*. We may conclude that sequences of different specimens of a species clustered together with the maximum support or nearly so, while cases of tight clustering of different species corresponded to the notion of their relatedness based on morphology.

It is noteworthy that the sequences of two specimens of *S. croceolum* from its West Siberian isolate (specimens Sc-41 and Sc-42) (Kosterin 2002) did not show divergence from those from the main Far Eastern range of the species (the rest specimens) but nested among them.

## Discussion

### The marker proposed

We may conclude that the phylogenetic information provided by the proposed marker well resolved the overall phylogenetic relationships of Odonata at taxonomic levels of families, genera and species (Fig. 2), with few notable exceptions, which could actually reflect weak points of the currently accepted taxonomic system (Dijkstra et al. 2013).

The high conservation of the histone H3 and H4 proteins is paralleled by the high conservation of their coding sequences, the variation of which is nearly confined to synonymous substitutions (Stein et al. 1984; Doenecke et al. 1997; Eirín-López et al. 2009). This allowed us to design primers highly specific to these particular genes but of a very broad specificity as to the biological objects. It is noteworthy that the substitutions in the histone coding sequence, the overwhelming majority of which are synonymous, in the histone H3-H4 region provide enough variation for satisfactory resolving phylogenetic relations between the studied species (Fig. 3).

Because of this, the histone H3 gene coding sequence has been broadly used as a phylogenetic marker for short evolutionary distances, e.g. in Odonata by Carle et al (2015), with conserved positions permitting universal primers whereas the phylogenetic signal was mostly provided by synonymous substitutions.

The here proposed marker, the histone H3-H4 region, has an advantage of possibility to design highly universal primers matching the most conserved eukaryotic coding regions, those of histones H3 and H4. Note, that we used LH4_2R primer designed to match the sequence of a lepidopteran, *B. mori*. The amplified fragment contains most of the coding region of histone H4 and about a half of that of histone H3, and ca 250 b.p. long spacer between them. No significant adaptive constraint is expected for variation of the spacer, which hence has a neutral regime of evolution and may serve as molecolar clock.

The histone H3-H4 region involving both highly conserved sequences and a non-coding spacer plus being tandemly repeated makes the proposed phylogenetic marker similar in biological and technical respects to such popular nuclear marker as the ITS region of the nucleolus organiser (for its use in Odonata see Hovmöller & Johansson 2004; Dumont et al. 2010; Schneider et al. 2023) including the internal spacers *ITS1* and *ITS2*. The length of ca 250 b.p. of the spacer in the here proposed marker is comparable to ca 200 b.p. of *ITS1* and ca 160 b.p. of *ITS2* (these figures are for Odonata). Both markers, the ITS and histone H3-H4 region, have comparable lengths (ca 900 b.p.) and are encoded by the nuclear genome but functionally unrelated. Hence the histone H3-H4 region can be used for the same purposes as the ITS region. Moreover, comparison of phylogenetic resolution of Odonata at the family level (Figs 2, 5) showed that the H3-H4 region adequately resolved the phylogeny of Odonata at the family level (Fig 2) while the ITS region rather failed to do this (Fig. 5), so the use of the former is preferable at the family level.

Therefore, the use of histone H3-H4 region can update the traditional analysis of the ITS region with about the same amount of independent phylogenetic information of the same nature. A joint analysis of both similar but unrelated nuclear markers, ITS and H3-H4 regions (by their concatenation or, better, involving software specially designed for simultaneous analysis of different markers), is expected to provide a more robust phylogenetic inference than the analysis of ITS alone. Judging by the phylogenetic trees obtained (Figs 2, 6), the use of the histone H3-H4 region as a phylogenetic marker is highly recommendable at the levels of species and genera. Since it correctly resolves the family structure of the order, with few exceptions, it could be also useful at the level of families as well, but better as an additional marker analysed together with other phylogenetic markers.

Because of conservativeness of the histone H3 and H4 genes, the new marker can be used with the primers provided herein for any Odonata and those insects with the same order and orientation of the histone H3 and H4 genes in the histone repeat, e.g. Diptera exemplified in the present study by such genera as *Aedes* Meigen, 1818 and *Anopheles* Meigen, 1818, used as outgroup in the phylogenetic trees of Figs 2-4. The same order was found in *D. melanogaster* (Goodenough 1984), which represents another suborder of Diptera, as well as in the *Formica* Linnaeus, 1758 ants. For the use of the here proposed marker in insects with other order or orientations of these genes, relevant other primers have to be worked out. For example, in Lepidoptera, where the genes of histones H3 and H4 have parallel orientation, the same LH4-2R primer can be used in combination with a primer of the sequence which is reverse complement of that of the Hex-AR primer. In this case, the portion of the coding sequence of the H3 gene will be smaller, 37 bp, and the spacer will be somewhat longer - about 900 - 1400 bp.

### Sympetrum spp

The analysis of the new marker at once yielded two rather unexpected results, both concerning the species *S. croceolum* and *S. uniforme* (Fig. 6). First, *S. uniforme* appeared to be an inner branch nested inside *S. croceolum*. This result appeared robust regardless of the methods and models of phylogenetic reconstructions (not shown). These species are obviously related but well distinguishable by the morphology of the male genitalia and female vulvar scale, the wing coloration (dull, complete but gradually changing in intensity in *S. uniforme* versus bright but with gaps in *S. croceolum*) and size (the former is somewhat larger). In East Asia, they usually co-exist in a wide range of lentic habitats (Onishko & Kosterin 2022), while *S. croceolum* also has an isolated range fragment in the southern West Siberia (Kosterin 1987; 2002; Popova & Haritonov 2020). It should be stressed that specimens of *S. uniforme*, identified by external characters, formed a highly supported cluster (Fig. 6). There is no doubt that *S. croceolum* and *S. uniforme* are *bona species*. The phylogenetic pattern obtained, where the sequences of *S. uniforme* are nested inside those of *S. croceolum*, suggests *S. uniforme* being a phyletic descendant of *S. croceolum*, which hence appeared paraphyletic. This contradicts the cladistic approach in systematics and the phylogenetic concept of species. At the same time, this pattern fits well the so-called punctuated equilibria mode of speciation (Eldredge & Gould 1972), suggesting that speciation takes place for short time periods in evolutionary scale (tens to hundreds of thousand years) in small, isolated populations in the periphery of parental species’ ranges, while species exist almost unchanged for millions of years (evolutionary stasis). This concept better fits basics of evolutionary genetics (Mayr 1963; Berdnikov 1999) than the earlier prevailing model of gradual divergent evolution. In the punctualistic point of view, species ‘propagate’ as if being individuals, with younger species often co-existing with their parental species.

In the phylogenetic tree based on the histone H3-H4 region (Fig. 2), two analyzed specimens of *S. croceolum* from its West Siberian isolate (specimens Sc-41 and Sc-42 from Lake Manzherok) lacked supported divergence from specimens from the main Far Eastern range of the species (the rest specimens),which is quite remarkable. Specimens from the West Siberian isolate (Kosterin 2002) differ from the Far Eastern specimens by much more developed wing amber colour and appearance of a brown enfumation in the wing apical parts and so were for a long time supposed to represent a separate subspecies (Kosterin 1987; Popova & Haritonov 2020), which, however, has not got a name yet. This lack of divergence at molecular level suggests the West Siberian isolate to be very young in evolutionary time scale and well fits its hypothetic Holocene age, implying the range split after Atlantic time (Kosterin 2002), as well as in some nemoral species of Lepidoptera (Dubatolov & Kosterin 2000; Solovyev et al. 2015; 2022). This, however, does not exclude a subspecies rank of the West Siberian population(s), since subspecies are entities of well-defined geographical variation for some phenotype characters and so imply specific divergence of the genes determining these characters rather than of the entire genome.

In all phylogenetic reconstructions from sequences of the histone H3-H4 region (Figs 2, 6), *S. fonscolombii* is strongly diverged from the rest of the genus *Sympetrum*. The same result was earlier obtained by Pilgrim & von Dohlen (2012) who undertook a molecular phylogenetic study of *Sympetrum* and related genera based on the joint analysis of the nuclear marker *EF-1α* and ITS2, and the mitochondrial genes *16S, tRNAvaline, 12S*, and *CO*I. This divergent position of *S. fonscolombii* has long ago been recognised at the level of phenotype, resulting in a suggestion to move *S. fonscolombii* to the genus *Tarnetrum* Needham & Fischer, 1936 (Schmidt 1987). This genus was erected for two Nearctic species, *Mesothemis corrupta* Hagen 1861, and *M. illota* Hagen, 1861 (mentioned therein as well as presently considered in combinations *Sympetrum illotum* and *S. corruptum*), with *M. illota* (sub. *S. illotum*) indicated as the type species (Needham & Fischer 1936). Of them, *S. corruptum* is quite closely related to *S. fonscolombii* (together with *S. villosum* Ris, 1911 and *Nesogonia blackburni* (McLachlan, 1883)) while *S. illotum* is not (Pilgrim & von Dohlen 2012). Since the type species of the genus *Tarnetrum* is not closely related to *S. fonscolombii*, this genus is not suitable for the latter species (Dijkstra & Kalkman 2015). Therefore, the genus *Sympetrum* in the current sense deserves further reconsideration, maybe with erection of a new genus at least for the *fonscolombii*-group *sensu* Pilgrim et al. (2012).

## Acknowledgements

The work was supported by the project FWNR-2022-0019 of the Institute of Cytology and Genetics SB RAS. DNA sequencing was performed at the SB RAS Genomics Core Facility, Novosibirsk. Assembly of sequences from SRA archive was performed at Siberian SuperComputer Computational Facility, Novosibirsk.

## References

Avise J.C. 2000. Phylogeography: the history and formation of species. Harvard University Press, Cambridge

Avise J.C. 2009. Phylogeography: retrospect and prospect. Journal of Biogeography 36: 3–15 10.1111/j.1365-2699.2008.02032.x

Ballard J.W.O. & Whitlock M.C. 2004, The incomplete natural history of mitochondria. Molecular Ecology 13: 729–744 10.1046/j.1365-294x.2003.02063.x

Berdnikov V.A. 1999. Evolution and progress. Pensoft, Sofia-Moscow

Carle F.L., Kjer K.M. & May M.L. 2015. A molecular phylogeny and classification of Anisoptera. Arthropod Systematic and Phylogeny 73 (2): 281–301 10.3897/asp.73.e31805

Bybee S.M., Kalkman V.J., Erickson R.J., Frandsen P.B., Breinholt J.W., Suvorov A., Dijkstra K.B., Cordero-Rivera A., Skevington J.H., Abbott J.C., Sanchez Herrera M., Lemmon A.R., Moriarty Lemmon E., Ware J.L. 2021 Phylogeny and classification of Odonata using targeted genomics. Molecular Phylogenetics and Evolution 160: 107115. 10.1016/j.ympev.2021.107115.

Cheng Y.-C., Chen M.-Y., Wang J.-F., Liang A.P. & Lin C.-P. 2024. Some mitochondrial genes perform better for damselfly phylogenetics: species- and population-level analyses of four complete mitogenomes of *Euphaea* sibling species. Systematic Entomology 43 (4): 702–715 10.3389/10.1111/syen.12299

Cheng Z, Li Q, Deng J, Liu Q & Huang X. 2023. The devil is in the details: Problems in DNA barcoding practices indicated by systematic evaluation of insect barcodes. Frontiers in Ecology and Evolution 11: 1149839 10.3389/fevo.2023.1149839

Chevreux B., Wetter T. & Suhai S. 1999. Genome sequence assembly using trace signals and additional sequence information. Computer Science and Biology: Proceedings of the German Conference on Bioinformatics (GCB) 99: 45–56

Deng J., Assandri G., Chauhan P., Futahashi R., Galimberti A., Hansson B., Lancaster L.T., Takahashi Y., Svensson E.I. & Douploy A. 2021. *Wolbachia*-driven selective sweep in a range expanding insect species. BMC Ecology and Evolution 21: 181 10.1186/s12862-021-01906-6

Dijkstra K.D.B., Bechly G., Bybee S.M., Dow R.A., Dumont H.J., Fleck G., Garrison R, Hämäläinen M., Kalkman V.J., Karube H., May M.L., Orr A.G., Paulson D.R., Rehn A.C., Theischinger G., Trueman J.W.H., Van Tol J., Von Ellenrieder N. & Ware J. 2013. The classification and diversity of dragonflies and damselflies (Odonata). In: Zhang, Z.-Q.(Ed.) Animal Biodiversity: An Outline of Higher-level Classification and Survey of Taxonomic Richness (Addenda 2013). Zootaxa 3703 (1): 36–45. 10.11646/zootaxa.3703.1.9

Dijkstra J.P. & Kalkman V. 2015. Taxonomy. In: Boudot, J.P. & Kalkman, V.J. (eds), Atlas of the European Dragonflies and Damselflies. KNNNV Publishing, the Netherlands, p. 15–25

Doenecke D., Albig V., Bode C., Drabent B., Franke K., Gavenis K. & Witt O. 1997. Histones: genetic diversity and tissue-specific gene expression. Histochemistry and Cell Biology 107: 1–10 10.1007/s004180050083

Dow R.A.; Butler S.G.; Reels G.T.; Steinhoff O.M.; Stokvis F. & Unggang L. 2019. Previously unpublished Odonata records from Sarawak, Borneo, part IV: Bintulu Division including the Planted Forest Project and Similajau National Park. Faunistic Studies in south-eastern and Pacific Island Odonata 27: 1–66

Dubatolov V.V. & Kosterin O.E. 2000. Nemoral species of Lepidoptera (Insecta) in Siberia: a novel view on their history and the timing of their range disjunctions. Entomologica Fennica 11(3): 141–166 10.33338/ef.84061

Dumont H.J., Vierstraete A. & Vanfleteren J.R. 2010. Molecular phylogeny of the Odonata (Insecta). Systematic Entomology 35: 6–18 10.1111/j.1365-3113.2009.00489.x

Eirín-López J.M., González-Romero R., Dryhurst D., Méndez J. & Ausio J. 2009. Long-term evolution of histone families: old notions and new insights into their mechanisms of diversification across eukaryotes. In: P. Pontarotti (ed.) Evolutionary Biology: Concept, Modeling, and Application. Heidelberg, Springer, p 139–162

Eldredge N. & Gould S.L. 1972. Punctuated equilibria: an alternative to phyletic gradualism. In: Schopf, T.J.M. (ed.) Models in palaeobiology. Freeman, Cooper & Co., San Francisco, p. 82–115

Ferreira, S., Boudot, J.-P., El Haissoufi, M., Alves, P.C., Thompson, D.J. & Watts, P.C. (2016) Genetic distinctiveness of the damselfly *Coenagrion puella* in North Africa: an overlooked and endangered taxon. Conservation Genetics 17: 985–991. 10.1007/s10592-016-0826-5

Ferreira S., Lorenzo-Carballa M.O., Torres-Cambas Y., Cordero-Rivera A., Thompson D.J. & Watts P.C. 2014. New EPIC nuclear DNA sequence markers to improve the resolution of phylogeographic studies of cenagrionids and other odonates. International Journal of Odonatology 17 (2-3): 135–147 10.1080/13887890.2014.950698

Filip E. & Skuza L. 2021. Horizontal Gene Transfer Involving Chloroplasts. International Journal of Molecular Science 22 (9): 4484 10.3390/ijms22094484)

Futahashi R., Kawahara-Miki R., Kinoshita M., Yoshitake K., Yajima S., Arikawa K., Fukatsu T. 2015. Extraordinary diversity of visual opsin genes in dragonflies. Proceedings of the National Academy of Sciences of the U. S. A. 112 (11): E1247–56. 10.1073/pnas.1424670112

Galimberti A.; Assandri G.; Maggioni D.; Ramazotti F.; Baroni D.; Bazzi G.; Chiandetti I.; Corso A.; Ferri V.; Galuppi M.; Ilahiani L.; La Porta G.; Laddaga L.; Landi F.; Mastropasqua F.; Ramellini S.; Santinelli R.; Soldato G.; Surdo S. & Casiraghi M. 2021. Italian odonates in the Pandora’s box: a comprehensive DNA barcoding inventory shows taxonomic warnings at the Holarctic scale. Molecular Ecology Resources 21: 183–200 10.1111/1755-0998.13235

Geiger M.; Koblmüller S.; Assandri G.; Chovanec A.; Ekrem T.; Fischer I.; Galimberti A.; Grabowski M.; Haring E.; Hausmann A.; Hendrich L.; Koch S.; Mamos T.; Rothe U.; Rulik B.; Rewicz T.; Sittenthaler M.; Stur E.; Tończyk G.; Zangl L. & Moriniere J. 2021. Coverage and quality of DNA barcode references for Central and Northern European Odonata. PeerJ 9: e11192 10.7717/peerj.11192

Goodenough U. 1984. Genetics. Third Edition, CBS College Publishing, Hong Kong

Gurdon C., Svab Z., Feng Y., Kumar D. & Maliga P. 2016. Cell-to-cell movement of mitochondria in plants. Proceedings of the National Academy of Sciences of the U. S. A. 113: 3395–3400 10.1073/pnas.1518644113)

Hovmöller R. & Johansson F. 2004. A phylogenetic perspective on larval spine morphology in Leucorrhinia (Odonata: Libellulidae) based on ITS1, 5.8s and ITS2 rDNA sequences. Molecular Phylogenetics and Evolution 30: 653–662 10.1016/S1055-7903(03)00226-4

Karube H., Futahashi R., Sasamoto A. & Kawashima I. 2011. Taxonomic revision of Japanese Odonata species, based on nuclear and and mitochondrial gene genealogies and morphological comparison with allied species. Part I. Tombo 54: 75–106

Kohli M., Letsch H., Greve C., Bethoux O., Deregnaucourt I., Liu S., Zhou X., Donath A., Mayer C., Podsiadlowski L., Gunkel S., Machida R., Niehuis O., Rust J., Wappler T., Yu X., Misof B., Ware J. 2021. Evolutionary history and divergence times of Odonata (dragonflies and damselflies) revealed through transcriptomics. iScience 24(11):1 03324. 10.1016/j.isci.2021.103324

Kosterin O.E. 1987. Discovery of East-Asiatic dragonfly (Odonata, Libellulidae) at the Manzherock Lake (Altay). In Cherepanov A.I. (ed.) [Insects, Mites, and Helmints. New and little-known species of the fauna of Siberia]. Nauka (Siberian Division), Novosibirsk, p. 57–63 [in Russian, English summary]

Kosterin O.E. 2002. Western range limits and isolates of eastern odonate species in Siberia and their putative origins. Odonatologica 34(3): 219–242

Larkin M.A., Blackshields G., Brown N.P., Chenna R., McGettigan P.A., McWilliam H., Valentin F., Wallace I.M., Wilm A., Lopez R., Thompson J.D., Gibson T.J. & Higgins D.G. 2007. Clustal W and Clustal X version 2.0. Bioinformatics 23: 2947–2948 10.1093/bioinformatics/btm404

Lorenzo-Carballa M.O., Sanmartín-Villar I. & Cordero-Rivera A. 2022. Molecular and morphological analyses support different taxonomic units for Asian and Australo-Pacific forms of *Ischnura aurora* (Odonata, Coenagrionidae). Diversity 14 (8): 606. 10.3390/d14080606

Lowe C.D., Harvey I.F., Thompson D.J. & Watts P. C. 2008. Strong genetic divergence indicates that congeneric damselflies *Coenagrion puella* and *C. pulchellum* (Odonata: Zygoptera: Coenagrionidae) do not hybridise. Hydrobiologia 605, 55–63. 10.1007/s10750-008-9300-9

Mayr E. 1963. Zoological species and evolution. The Belknap Press of Harward University Press, Cambridge, Massachusetts Needham J.G. & Fischer E. 1936. The nymphs of North American libelluline dragonflies. Transactions of the American Entomological Society 62 (2): 107–116.

Onishko V. & Kosterin O. 2021. Dragonflies of Russia. Illustrated Photo Guide. Phyton XXI. Moscow [in Russian, English abstract]

Ožana S., Dolný A. & Pánek T. 2022. Nuclear copies of mitochondrial DNA as a potential problem for phylogenetic and population genetic studies of Odonata. Systematic Entomology 47 (4): 591– 602. 10.3389/10.1111/syen.12550

Pilgrim E.M. & von Dohlen C.D. 2012. Phylogeny of the dragonfly genus *Sympetrum*. Organisms Diversity & Evolution 12: 281–291 10.1007/s13127-012-0081-7

Popova O.N. & Haritonov A.Y. 2020. On the distribution of *Sympetrum croceolum* in the Russian part of its range (Odonata: Libellulidae). Odonatologica 49 (1/2): 29–49 10.5281/zenodo.3823325

Schmidt E. 1983. Generic reclassification of some Westpalearctic Odonata taxa in view of their Nearctic affinities. Advances in Odonatology 3: 135–145

Schneider T., Vierstraete A., Kosterin O.E, Ikemeyer D., Hu F.-S., Snegovaya N. & Dumont H.J. 2023. Molecular phylogeny of Holarctic Aeshnidae with a focus on the West Palaearctic and some remarks on its genera worldwide (Aeshnidae, Odonata). Diversity 15: 950 10.3390/d15090950

Solovyev V.I., Bogdanova V.S., Dubatolov V.V. & Kosterin O.E. 2015. Range of a Palearctic uraniid moth *Eversmannia exornata* (Lepidoptera: Uraniidae: Epipleminae) was split in the Holocene, as evaluated using histone H1 and COI genes with reference to the Beringian disjunction in the genus *Oreta* (Lepidoptera: Drepanidae). Organisms Diversity & Evolution 15(2): 285–300 10.1007/s13127-014-0195-1

Solovyev V.I., Dubatolov V.V., Vavilova V.Y. & Kosterin O.E. 2022. Estimating range disjunction time of the Palaearctic Admirals (Limenitis L.) with COI and histone H1 genes. Organisms Diversity & Evolution 22: 975–1002 10.1007/s13127-022-00565-9

Staden R., Judge D.P., Bonfield J.K. 2003. Managing sequencing projects in the GAP4 environment. In: Krawetz S.A. & Womble D.D. (eds) Introduction to Bioinformatics. A Theoretical and Practical Approach. Eds. Human Press Inc., Totawa

Stein G.S., Stein J.L. & Marzluff W.F. (Eds.). 1984. Histone Genes: Structure, Organization, and Regulation. John Willey & Sons, New York

Tamura K., Stecher G., Peterson D. & Kumar S. 2013. MEGA6: molecular evolutionary genetics analysis version 6.0. Molecular Biology and Evolution 30: 2725–2729 10.1093/molbev/mst197

